## Supplemental for "Maf/ham1-like pyrophosphatases of non-canonical nucleotides are host-specific partners of viral RNA-dependent RNA polymerases"

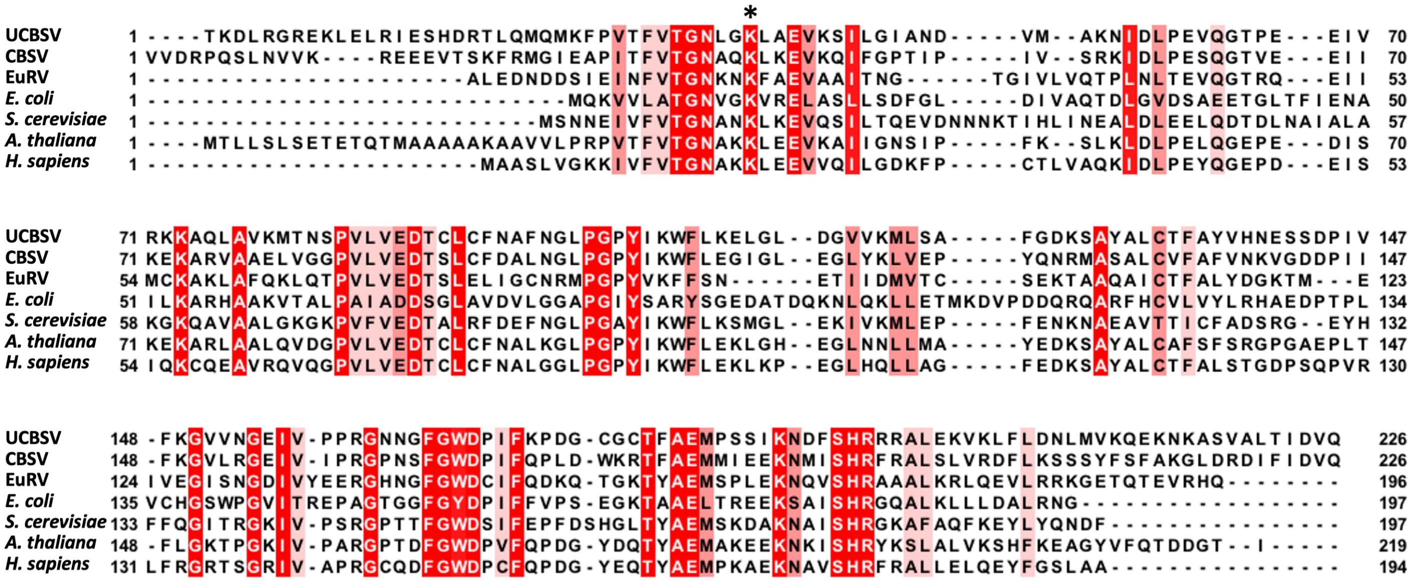

Supplementary Figure S1

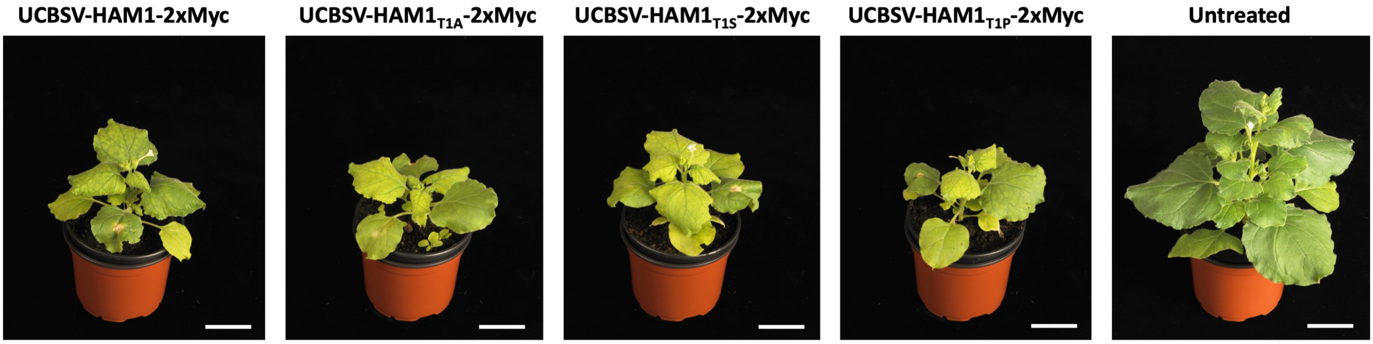

Supplementary Figure S2

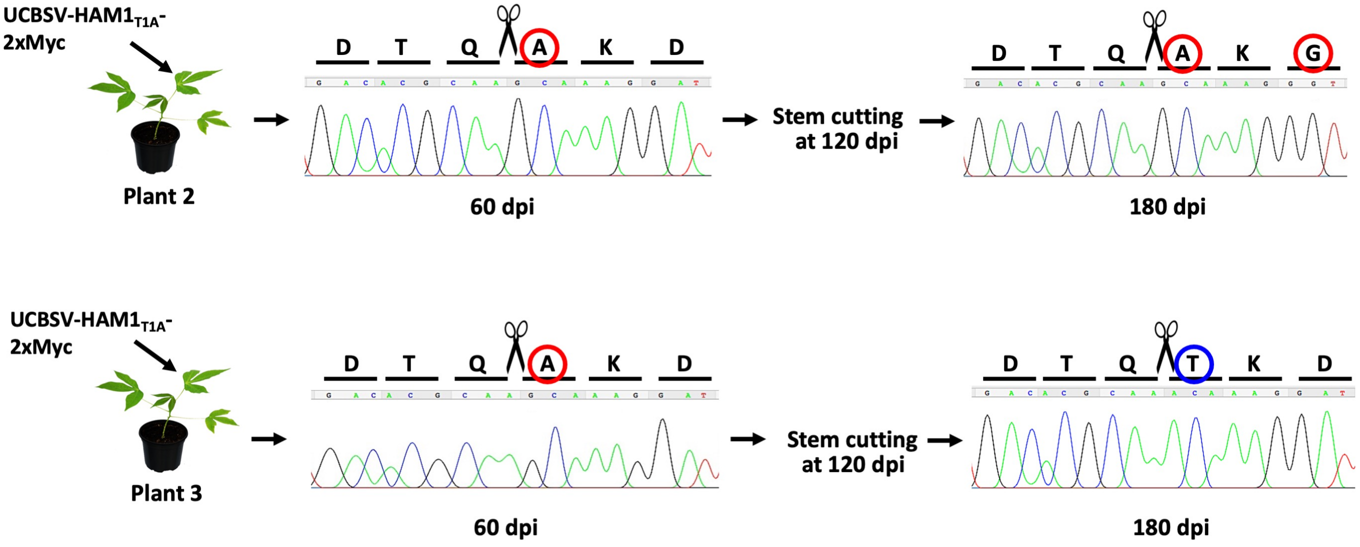

Supplementary Figure S3

**Legends for Supplementary Figures**

**Supplementary Figure 1. Alignment of HAM1 proteins from the indicated potyvirids and organisms.** The intensity of red colour correlates with the degree of amino acid identities. Black asterisk indicates the fully-conserved K present in all known HAM1 proteins that was mutated by A in this study (see Figure 3).

**Supplementary Figure 2. Infection symptoms in *Nicotiana benthamiana* produced by 2xMyc-tagged UCBSV mutated in the first amino acid of HAM1.** Representative pictures of infected and non-treated *N. benthamiana plants* taken at 12 days post-inoculation. Bar = 4 cm. This image complements the information shown in Figure 5.

**Supplementary Figure 3. Experimental evolution of 2xMyc-tagged UCBSV that carries the T1A mutation in plants multiplied by stem cutting.** Chromatograms of Sanger sequencing results of the DNA fragment of interest amplified by RT-PCR. RNA samples obtained from upper non-inoculated leaves of cassava plants inoculated with the UCBSV-HAM1$_{T1A}$-2xMyc mutant, and from plants derived from a stem cutting passage were used as template. Leaves for RNA preparation were harvested at 60- and 180-days post-infection (dpi). Residues derived from the original mutation and from the spontaneous second mutation are surrounded by a red circle. The amino acid that appears as the reversion to the wild type variant is surrounded by a blue circle.

**Supplementary table S1**. Oligonucleotides used in this study.

| Primer | Sequence (5'-3') |
|---|---|
| #3071 | ATGGGCTACAGTAAACGACAACGTCTTAAG |
| #3072 | ATGTGTTGAAAAGCATGCACTTGC |
| #3123 | ATGAATGGTGATGATTTGATCATAGCTATTAACC |
| #3124 | ATGTGACTTGTTCCTCGCCATC |
| #3127 | ATGGCGAAGCACAAGTATAACAGAGATAAG |
| #3128 | AAATGCTTGGAAGATATTGGTCTCG |
| #3129 | ATGAATGGGGATGATCTGATCCTCAATG |
| #3130 | AAGCTGACTGTGATACCTTTGGAGG |
| #3160 | AGATATCAATGAAGAATGGTTGATAGG |
| #3162 | ACCGTTCAAGTCTTCCTCGGAGATTAGCTTTTGTTCACCGTTAATTA ACCCAGCCACACTTGCTTTATTCTTCTC |
| #3163 | CCTGAACGTCAATCGTTAGAGCAAGATCCTCTTCTGAAATTAATTT TTGTTCACCGTTCAAGTCTTCCTCGG |
| #3205 | CAGGAAATTTGGGAGCATTAGCAGAGGTGAAGTCTATTCTTG |
| #3206 | CACCTCTGCTAATGCTCCCAAATTTCCTGTCACAAATG |
| #3255 | TACCTGCAGATGTCGACTATC |
| #3256 | CAAAGGTAGAAGCAGAAACTTACCTGGCTTTCCGTGCATAC |
| #3257 | GTAAGTTTCTGCTTCTACCTTTG |
| #3258 | CTGCATATCAACAAATTTTGG |
| #3259 | CCAAAATTTGTTGATATGCAGGTTTTACAAATTTGTGGATCAC |
| #3260 | CAGCAGCACACCACTTTAG |
| #3261 | GACACGCAAGCAAAGGATTTGAGAGGAAGA |
| #3262 | CAAATCCTTTGCTTGCGTGTCAACATAAC |
| #3263 | GACACGCAATCAAAGGATTTGAGAGGAAGA |
| #3264 | CAAATCCTTTGATTGCGTGTCAACATAAC |
| #3265 | GACACGCAACCAAAGGATTTGAGAGGAAGA |
| #3266 | CAAATCCTTTGGTTGCGTGTCAACATAAC |
| #3312 | CGAACATTGTTATGTTGACACGCAAG |
| #3313 | CTTGCGTGTCAACATAACAATGTT |
| #3358 | GCTCCTCGCCCTTGCTCACCATGGCCTGAACGTCAATCGTTAGAGC |
| #3360 | ATGGTGAGCAAGGGCGAGGAGC |
| #3361 | CTTGCACATCGATTGTCAAGGTACCCTTGTACAGCTCGTCC |
| #3647 | ATGGCAAAGCATAAGTATAATCGGG |
| #3648 | ACGCCGGTTTACTCAAAATGC |
| #3649 | ATGAATGGTGATGATCTTATATTGAATGC |
| #3650 | AGCTCTTTCGAAATTCTAGAGCG |

**Supplementary table S2.** Templates and name of primers used for PCR amplifications during the construction of the indicated plasmids are shown.

| Plasmid | Mutant | PCR | Primer Forward* | Primer Reverse* | Template |
|---|---|---|---|---|---|
| pLX-UCBSVi | K38A | 1 | #3205 | #3130 | pLX-UCBSVi |
| | | 2 | #3160 | #3206 | pLX-UCBSVi |
| | | overlapped | #3160 | #3130 | PCR1 + PCR2 |
| pLX-UCBSVi-HAM1-2xMyc | T1A | 1 | #3261 | #3130 | pLX-UCBSVi-HAM1-2xMyc |
| | | 2 | #3160 | #3262 | pLX-UCBSVi-HAM1-2xMyc |
| | | overlapped | #3160 | #3130 | PCR1 + PCR2 |
| | T1S | 1 | #3263 | #3130 | pLX-UCBSVi-HAM1-2xMyc |
| | | 2 | #3160 | #3264 | pLX-UCBSVi-HAM1-2xMyc |
| | | overlapped | #3160 | #3130 | PCR1 + PCR2 |
| | T1P | 1 | #3265 | #3130 | pLX-UCBSVi-HAM1-2xMyc |
| | | 2 | #3160 | #3266 | pLX-UCBSVi-HAM1-2xMyc |
| | | overlapped | #3160 | #3130 | PCR1 + PCR2 |

*The sequences of oligonucleotides used in this study are shown in Supplementary table S1

**Supplementary table S3.** Templates and name of primers used for PCR amplifications during the construction of the indicated plasmids are shown.

| Plasmid | Virus | Fragment | Primer Forward[1] | Primer Reverse[1] | Template |
|---|---|---|---|---|---|
| pENTR1A | EuRV | NIa | #3071 | #3072 | cDNA from an infected plant[2] |
| | | NIb$_C$-HAM1-CP$_N$ | #3123 | #3124 | cDNA from an infected plant[2] |
| | UCBSV | NIa | #3127 | #3128 | pLX-UCBSVi |
| | | NIb$_C$-HAM1-CP$_N$ | #3129 | #3130 | pLX-UCBSVi |
| | CBSV | NIa | #3647 | #3648 | pYES2-CBSV-F2[3] |
| | | NIb$_C$-HAM1-CP$_N$ | #3649 | #3650 | pYES2-CBSV-F2[3] |

[1]The sequences of oligonucleotides used in this study are shown in Supplementary table S1

[2]cDNA was prepared from RNA of *Euphorbia milii* leaves infected with EuRV (Leibniz Institute DSMZ - German Collection of Microorganisms and Cell Cultures GmbH)

[3]This plasmid harbours a cDNA segment that corresponds to the 3'half part of the CBSV genome (unpublished). Kindly provided by Gary Foster.
